# Simultaneous Knockouts of the Oxytocin and Vasopressin 1b Receptors in Hippocampal CA2 Impair Social Memory

**DOI:** 10.1101/2023.01.30.526271

**Authors:** Adi Cymerblit-Sabba, Caroline Walsh, Kai-Zheng Duan, June Song, Oliver Holmes, W. Scott Young

## Abstract

Oxytocin (Oxt) and vasopressin (Avp) are two neuropeptides with many central actions related to social cognition. The oxytocin (Oxtr) and vasopressin 1b (Avpr1b) receptors are co-expressed in the pyramidal neurons of the hippocampal subfield CA2 and are known to play a critical role in social memory formation. How the neuropeptides perform this function in this region is not fully understood. Here, we report the behavioral effects of a life-long conditional removal (knockout, KO) of either the Oxtr alone or both Avpr1b and Oxtr from the pyramidal neurons of CA2 as well as the resultant changes in synaptic transmission within the different fields of the hippocampus. Surprisingly, the removal of both receptors results in mice that are unable to habituate to a familiar female presented for short duration over short intervals but are able to recognize and discriminate females when presented for a longer duration over a longer interval. Importantly, these double KO mice were unable to discriminate between a male littermate and a novel male. Synaptic transmission between CA3 and CA2 is enhanced in these mice, suggesting a compensatory mechanism is activated to make up for the loss of the receptors. Overall, our results demonstrate that co-expression of the receptors in CA2 is necessary to allow intact social memory processing.

## Introduction

Social memory is critical for the structure and stability of relationships and hierarchies in societies across species. Deficits in social memory are commonly associated with neuropsychiatric disorders^1^. While the involvement of the hippocampus in acquiring and retrieving social memories is established, there is much to learn yet about the underlying mechanisms. Of special interest is the CA2 hippocampal subfield which plays a critical role in acquisition, consolidation and retrieval of social memory^2–4^.

The pyramidal neurons in CA2 co-express the vasopressin (Avp) 1b and oxytocin (Oxt) receptors (Avpr1b and Oxtr, respectively), through which the important neuropeptides, Avp and Oxt, carry their modulatory action on social cognition and social behavior^5–7^. Previously, we showed that loss of the Avpr1b from the pyramidal neurons of the CA2 results in inability of mice to recognize a familiar conspecific^6–7^. We also described an enhancement of social recognition following activation of the paraventricular nucleus (PVN) terminals in CA2, mediated by the Avpr1b^2^ and, further, we were able to induce atypical partner preference behavior in mice through similar activation of the same neuronal pathway^8^.

Oxt release and binding within the hippocampus have been linked to social behaviors and memory^9^. Yet, it’s role in social memory modulation particularly in the pyramidal neurons of the CA2 subfield remains unclear. To the best of our knowledge, two publications addressed the question of social memory effects following Oxtr removal from the hippocampus. Both describe virally targeted conditional knockout of Oxtr in CA2/CA3 hippocampal areas of adult mice and result in different behavioral observations. In the first study^10^, group housed mice were repeatedly presented (trial 1-3) with a male mouse (C57BI/6J) for 5 minutes with an interval of 5 minutes between trials before attempted discrimination between the familiar and a novel male (trial 4). Both recognition and discrimination were impaired. In the second study^11^, group housed mice were presented with a novel juvenile male mouse (trial 1) for 5 minutes then required to discriminate between the familiar and a novel juvenile (trial 2) following either 10 minute, 24 hours or 7 days interval. No impairments were observed in discrimination following short intervals (10 minutes or 24 hours) but only following the longer 7 days interval. In a separate paradigm, mice were isolated for 7 days then tested in habituation dishabituation test with CD1 ovariectomized females. Mice were repeatedly presented with the same female (trails 1-4) for 1 minute with a 10 minute interval then with a novel one (trial 5). No impairments were observed.

In humans, intranasal Avp administration before encoding enhances the feeling of familiarity with images of positive and negative faces compared to neutral faces^12^ and in males, enhances the recognition of sexual cues in images^13^. Intranasal administration of Oxt post learning enhances immediate (30 minutes) and delayed (24 hours) recognition of images with angry and neutral faces but not happy ones^14^. Numerous other studies have shown different effects^15^, demonstrating the importance of the context used. Similarly, social memory performance in rodents depends on various factors, including the species, sex, age and emotional status of the animal during the investigation^16^. These factors, as well as reproducibility, should be considered when drawing conclusions regarding behavioral phenotypes.

Synaptic potentiation is induced in CA2 pyramidal neurons by Avpr1b and Oxtr agonists in slices of rat and mouse hippocampus with no significant effect on paired-pulse facilitation^17^. While the peptides appeared to be acting selectively on their respective receptor^17^, accumulated evidence suggest there is a significant cross-talk between the neuropeptides and the receptors^18^. Oxytocin can activate the vasopressin receptor and vasopressin can activate the oxytocin receptor and this cross-talk could potentially contribute to the regulation of the complex behaviors and physiological processes under their control.

To study the functional significance of the receptors’ co-expressions in CA2 pyramidal neurons, we selectively removed both. We observed impairments in social memory: mice were unable to habituate to a sexually naïve ovariectomized Balb/c female nor to recognize the novelty of another. They were also unable to discriminate between a littermate and a novel male mouse presented simultaneously. Surprisingly, they were able to recognize and discriminate sexually naïve ovariectomized Balb/c females in a different paradigm when allowed a longer time for familiarization. Also, synaptic transmission between CA3 and CA2 was enhanced in these mice, with reduced paired-pulse facilitation, suggesting a pre-synaptic compensatory mechanism.

Elucidating the mechanism by which the social neuromodulators Oxt and Avp function in CA2 is important for our understating of this hippocampal subregion’s functional role and may be of significant therapeutic potential in various psychiatric disorders.

## Methods

### Animals

All experiments were conducted according to US National Institutes of Health guidelines for animal research and were approved by the National Institute of Mental Health Animal Care and Use Committee. We bred our Avpr1b-Cre knockin mice^5^, in which Avpr1b is inactivated by the Cre recombinase coding sequence insertion and is expressed in CA2 pyramidal neurons, with mice containing loxP (f) sites flanking the *Oxtr* gene^19^. Homozygous *Oxtr*-floxed mice (*Oxtr^f/f^*) do not differ from WT littermates and express normal amounts of Oxtr^19^. To generate the experimental animals, C57Bl/6J mice with a confirmed genotype of fully floxed *Oxtr* and heterozygous for *Avpr1b* (*Oxtr^f/f^/ Avpr1b^+/Cre^*) were mated. The resulting offspring had the following genotypes: 1) *Oxtr^f/f^/Avpr1b^+/+^*; 2) *Oxtr^-/-^/Avpr1b^+/Cre^*; and 3) *Oxtr^-/-^/Avpr1b^Cre/Cre^*, and labeled as wildtype (WT), conditional *Oxtr* knockout with one functioning *Avpr1b* allele (HE), and conditional double knockout (DKO), respectively. This breeding regimen allow us to investigate the role of these receptors specifically in the CA2 pyramidal neurons where expression of *Oxtr* and *Avpr1b* overlap. All experimental mice were maintained on a C57Bl/6J background. Behavioral testing included 9 WT, 6 HE and 10 DKO. Stimulus mice used in social recognition testing were either sexually naïve, ovariectomized (OVX) Balb/c adult females or sexually-naïve C57Bl/6J males. All mice were maintained on a 12-h light cycle (lights off at 1500h) with *ad libitum* access to food and water and group housed except for the OVX Balb/c females which were housed individually.

### Behavioral testing

Behavioral tests were performed during the light cycle in a dimly lit room. Mice behaviors were recorded with Ethovision (Noldus Information Technology, Leesburg, VA) and coded by a trained observer, blind to the identity of the tested mouse.

### Social habituation-dishabituation (SHD)

Mice were brought to the testing room and habituated to a fresh mouse cage containing clean bedding for 30 min. An unfamiliar OVX Balb/C female was introduced repeatedly for 2 min with 5-min inter-trial intervals (ITI; T1-T4-stim 1). Then a novel OVX female was introduced (N-stim 2). After a 60-min ITI, the original female was reintroduced for 2 min (T5-stim 1). During ITIs, the female was returned to her home cage while the experimental animal remained in the testing cage. Mice were allowed to fully interact with anogenital and whole-body sniffing measured.

### Object habituation-dishabituation (OHD)

Mice were brought to the testing room and habituated to a fresh mouse cage containing clean bedding for 30 min. An unfamiliar object (Lego piece, Billund, Denmark, as stim 1) was introduced repeatedly for 2 min with 5-min ITIs (T1-T4-stim 1). Then a novel object was introduced (N-stim 2, 20ml scintillation vial with purple water as stim 2). After 60-min ITI, the original object was reintroduced for 2 min (T5-stim 1). Sniffing of the object, indicated by head movement and snout within 1 cm of the object, was measured.

### Social recognition-discrimination (SRD) test

Mice were brought to the testing room and habituated to a fresh mouse cage containing clean bedding for 30 min. Modified from Engelmann et al.^20^, an unfamiliar OVX female was introduced for 5 min (T1-stim 1), then reintroduced for 5 min after 30-min ITI (T2-recognition; stim 1) and after another 30-min ITI together with a novel female (T3/N-discrimination; stim1 and stim 2). Mice were allowed to fully interact with anogenital and whole-body sniffing measured. The discrimination index was calculated as (sniffing novel - sniffing familiar)/(sniffing novel + sniffing familiar).

### Long-term social recognition

Mice were brought to the testing room and habituated to a fresh mouse cage containing clean bedding for 30 min. An unfamiliar OVX female was introduced for 5 min (T1-stim 1), then reintroduced after 2 hours (T2-stim 1), 24 hours (T3-stim1) and 7 days (T4-stim1). Tested mice were placed back into their home cages during the ITIs. Mice were allowed to fully interact with anogenital and whole-body sniffing measured.

### Social novelty

Mice were brought to the testing room and habituated for 30 minutes. A test mouse was then placed in the center chamber of the Plexiglas three-chambered apparatus (each chamber 18×45×30cm) for a 5-min habituation with access to all chambers, followed by a 5-min test. In one side chamber, a male littermate was tethered with a small, beaded plastic zip tie. In the other outer chamber, a novel age- and weight-matched C57BL/6J male mouse was tethered. This prevented the stimulus mice from leaving their designated chambers but allowed their full access of the test animal (adapted from^21^). Stimulus mice were habituated to the collars for 15-30 minutes on a prior day. Anogenital and wholebody sniffing were measured.

### Slice electrophysiology

#### Slice preparation

Male mice (9–16 weeks) were anesthetized with isoflurane and decapitated. Brains were quickly removed and transverse hippocampal slices (350 μm) were prepared using a vibratome (Leica VT 1200S, Deer Park, IL) in ice-cold NMDG-HEPES solution containing (in mM): 93 NMDG (N-Methyl-D-glucamine diatrizoate), 2.5 KCl, 1.2 NaH_2_PO_4_, 30 NaHCO_3_, 20 HEPES, 25 D-glucose, 5 sodium ascorbate, 2 thiourea, 3 sodium pyruvate, 10 MgSO_4_ and 0.5 CaCl2 (adjusted to pH 7.3–7.4 with 10N HCl, bubbled with 95% O_2_/ 5% CO_2_). Brain slices were incubated in NMDG-HEPES solution (32°C) for 12-15 min, and then transferred to an immersed-type chamber and maintained in standard ACSF containing (in mM): 124 NaCl, 2.5 KCl, 1.2 NaH_2_PO_4_, 24 NaHCO_3_, 5 HEPES, 12.5 D-glucose, 2 MgSO_4_ and 2 CaCl_2_ (adjusted to pH 7.3–7.4, bubbled with 95% O_2_/ 5% CO_2_) at room temperature. The hippocampal slices were stabilized for 1–5 h before recording.

#### Extracellular field recordings

Field excitatory postsynaptic potentials (fEPSP) were recorded with a glass electrode (3~5MΩ, filled with ACSF) in a chamber allowing perfusion with oxygenized ACSF at 2.5 ml/min at 30°C. A bipolar stimulating electrode (impedance of 10 kΩ) was placed at a constant distance (150 μm approximately) in the stratum radiatum (SR) to stimulate Schaffer collateral fibers in the dorsal CA2 region or the stratum oriens (SO) to stimulate CA2 axons in the dorsal CA1 region. Spontaneous field potentials were recorded for 3 min in the SR or SO of dorsal CA2 region. Input–output (I–O) relations for field potentials were measured at the start of each experiment by applying a series of stimuli of increasing intensity with a 20 s interval. The protocol for the paired-pulse facilitation involved two stimulating pulses within a short (50 ms) inter-pulse interval. The paired-pulse ratio (PPR) was calculated as the ratio between the fEPSP amplitude evoked by the second stimulus divided by that induced by the first. Since PPR may vary with stimulus intensity, we adjusted stimuli to evoke initial fEPSPs of about 50% of maximal amplitude determined from input-output curves. Recording data were obtained using a Multiclamp 700B amplifier (San Jose, CA) and digitized using a Digidata 1550B board (San Jose, CA). Data were sampled at 10 kHz and analyzed with Clampfit 11.2 (San Jose, CA) and MATLAB R2021a (Chevy Chase, MD).

For the pharmacological experiments, acute transverse hippocampal slices were cut and transferred to an incubator filled with the standard ACSF saturated with 95% O_2_ and 5% CO_2_ (pH 7.4). The Avpr1b antagonist SSR149415 (1 μm at working concentration, Tocris, Cat# 6195), Oxtr antagonist L-371257 (1 μm at working concentration, Tocris, Cat# 2410), or the same volume of DMSO as control were introduced into the ASCF solution by a pipette above the bubbler. The slices were maintained in the incubator at room temperature for at least 1 h after the treatment, and then transferred to a recording chamber perfused with ACSF at 30 °C. The stimulus electrode was placed on the Schaffer collaterals to CA2, and the recording electrode was placed on the CA2 SR.

#### *In situ* hybridization for Oxtr and Avpr1b expression

Examination of expression of *Oxtr* and *Avpr1b* in the hippocampus was performed as previously described^22^. Male wildtype mice (N=3; *Oxtr^f/f^/Avpr1b^+/,+^*; WT), homozygous *Oxtr* and heterozygous *Avpr1b* (N=3; *Oxtr^-/-^/Avpr1b^+/Cre^*; HE), and simultaneous *Oxtr* and *Avpr1b* CA2 KO mice (N=3; *Oxtr^-/-^/Avpr1b^CreCre^*; DKO) were used. The Avpr1b (VB1-16301; transcript target NM_011924.2) and Oxtr (VB6-16868; target transcript NM_001081147) probes were designed and obtained from Affymetrix (Santa Clara, CA, USA), as was the ViewRNA duplex kit (Catalog number: QVT0013). Scans were obtained using a Zeiss Axio Scan Z1 (20x objective) and ZENlite software (Thornwood, NY, USA).

#### *In vitro* Oxtr Autoradiography

Sections containing the dorsal hippocampal CA2 area from five each of male WT, conditional *Oxtr* KO, and conditional DKO mice were examined for Oxtr binding as previously described^19^. As an additional control, we included two total Oxtr knockout mice^19^. Quantification was done with NIH Image J with integrated density (IntDen) within CA2 calculated after background subtraction outside of the hippocampus, thus: corrected total cell signal (CTCF)= IntDen (within CA2) – IntDen (outside hippocampus).

#### Statistical analysis

All data analyses were performed in Prism Software. For the comparison of two groups, two-tailed Student’s t-test or paired Student’s t-test was used for data that passed the normality and equal variance tests, and Mann-Whitney rank sum test was used for data that failed the normality and equal variances tests. For comparison of ≥ 3 groups with one factor, one-way ANOVA or one-way RM ANOVA was used for data that passed the normality and equal variance tests, and Kruskal-Wallis one-way ANOVA on ranks was used for data that failed the normality and equal variances tests. For comparison of ≥ 3 groups with two factors, two-way ANOVA was used. The assumptions of normality and homogeneity of variance were evaluated, and then two-way ANOVA analysis was performed. *P* < 0.05 was considered significant.

## Results

### *Oxtr* and *Avpr1b* expression in CA2 pyramidal neurons is greatly reduced in double knockout mice

We and others previously described the distribution of Oxtr and Avpr1b in the hippocampus^4, 11, 17, 23, 24^. To further understand the specific role of the receptors in social memory tasks in adulthood we generated new transgenic lines targeting either *Oxtr* or both *Oxtr* and *Avpr1b* in the pyramidal neurons of CA2 hippocampal subfield. We validated colocalization of *Oxtr* and *Avpr1b* transcripts in the CA2, with *Oxtr* extending into adjacent CA3 (Fig. 1a), consistent with previous reports^5, 23^. We also confirmed the reduction of *Oxtr* expression in the CA2 of HE and DKO mice and of *Avpr1b* expression in the DKO mice (Fig. 1b). Oxtr transcripts were greatly reduced in both conditional knockouts (HE and DKO), that we further confirmed with a binding experiment that demonstrated a significant decrease in the receptor expression in the region (Fig.1b-c). Remaining Oxtr receptors may reflect expression in the significant population of interneurons that express Oxtr^24^. A decrease in the transcripts of Avpr1b was also observed in both knockout lines, with the greatest loss in DKO mice (Fig. 1b, d). These results are consistent with the loss of one or both Avpr1b alleles in the HE or DKO mice, respectively. The chosen genetic approach allowed us to chronically remove the genes with anatomical specificity and thus gain insight into their neurodevelopmental role and associate possible behavioral phenotypes in adulthood.

**Figure 1.**
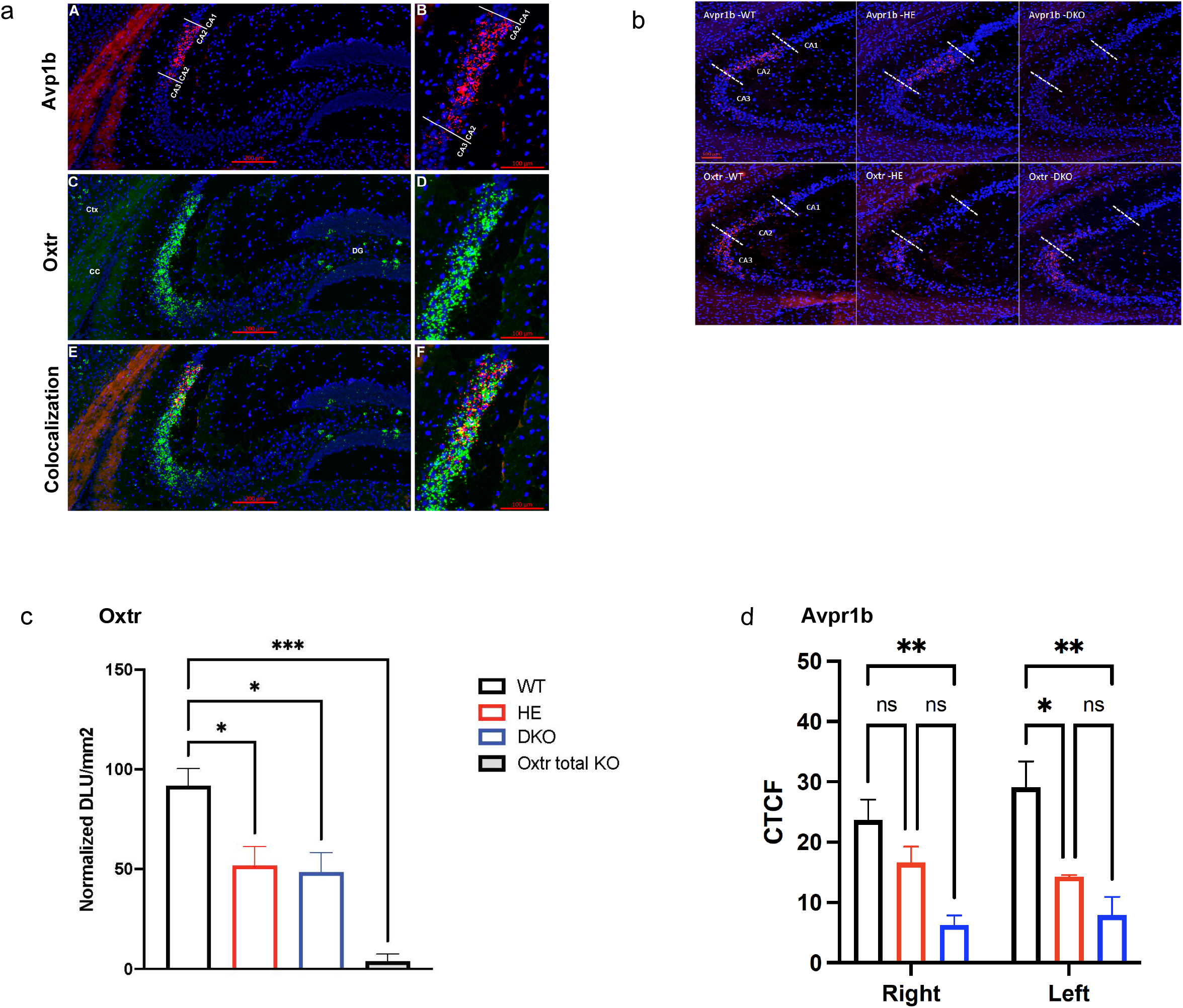
Validation of Oxtr and Avpr1b expression and removal from CA2 pyramidal neurons. a. A composite of Oxtr (green dots) and Avpr1b (red dots) gene expression identified by *in situ* hybridization in a section from the dorsal hippocampus of a wildtype mouse. The section is counterstained with DAPI in blue. Panels B, D and F are higher magnifications of the distributions in the dorsal CA2 subfield seen in A, C and E. Abbreviations: Ctx, neocortex; DG, dentate gyrus; cc, corpus callosum. b. A composite of Oxtr and Avpr1b expression in sections containing the dorsal CA2 area from the three different genotypes (WT: *Oxtr^f/f^/Avpr1b^+/+^*; HE: *Oxtr^-/-^/Avpr1b^+/Cre^*; and DKO: *Oxtr^-/-^/Avpr1b^Cre/Cre^*). The CA2 area is bounded by dashed lines. c. Coronal sections from adult mice of the three genotypes were used to evaluate Oxtr binding in the dorsal CA2 by *in vitro* receptor autoradiography. Oxtr binding is greatly reduced in the CA2 of both HE (*Oxtr^-/-^/Avpr1b^+/Cre^*) and DKO (*Oxtr^-/-^/Avpr1b^Cre/Cre^*) mice, shown by net grain intensities. One way ANOVA was significant for genotype F(3, 13)=10.02, P=0.0011, with Bonferroni multiple comparison further showing the difference between genotypes. d. Coronal sections from adult mice were used to evaluate transcripts expression of Avpr1b in the CA2. Corrected Total Cell Fluorescence, demonstrates the low presence of Avpr1b transcripts in the double knock out mice. Two way ANOVA (Genotype X Hemisphere) was significant for genotype F(2,5)=11.7, P=0.012, with within hemisphere Bonferroni multiple comparison showing a significant decrease. * P<0.05 **P<0.006***P<0.001. Data shown as mean± s.e.m.

### Conditional knockout of *Oxtr* or both *Oxtr* and *Avpr1b* in CA2 pyramidal neurons impairs social memory

The closely related neuropeptides Oxt and Avp have been extensively studied and are known to play a central role in social behaviors through their specific receptors in the brain. Yet, studies with different methodological approaches have resulted in varying behavioral observations^9^. Restricting the removal of the receptors to the pyramidal neurons of the CA2 allowed us to better delineate their function in social memory tasks with tissue-specific regulation. In rodents, novel conspecifics are met with increased interaction and sniffing that decrease with familiarization, as the animal forms a memory of the conspecific. We tested mice in four social memory tasks with varied exposure and interval times as well as different stimuli to find whether the life-long removal of the receptors interferes with their performance.

The loss of Oxtr (HE) from pyramidal neurons in CA2 impaired the ability of male mice to habituate to a familiar ovariectomized female as well as their ability to detect a novel one. Decreased sniffing was observed in the second trial (T2), which remained similar along the subsequent trials. This contrasts with the WT group that demonstrated decreased sniffing during the habituation as well as increased sniffing of the novel female in the dishabituation trial (N; Fig. 2a-b). The DKO mice, lacking both Oxtr and Avpr1b in the pyramidal neurons of CA2, showed no habituation across familiarization trials 1-4 and failed to detect the novelty of the female presented in the following trial (N).

**Figure 2.**
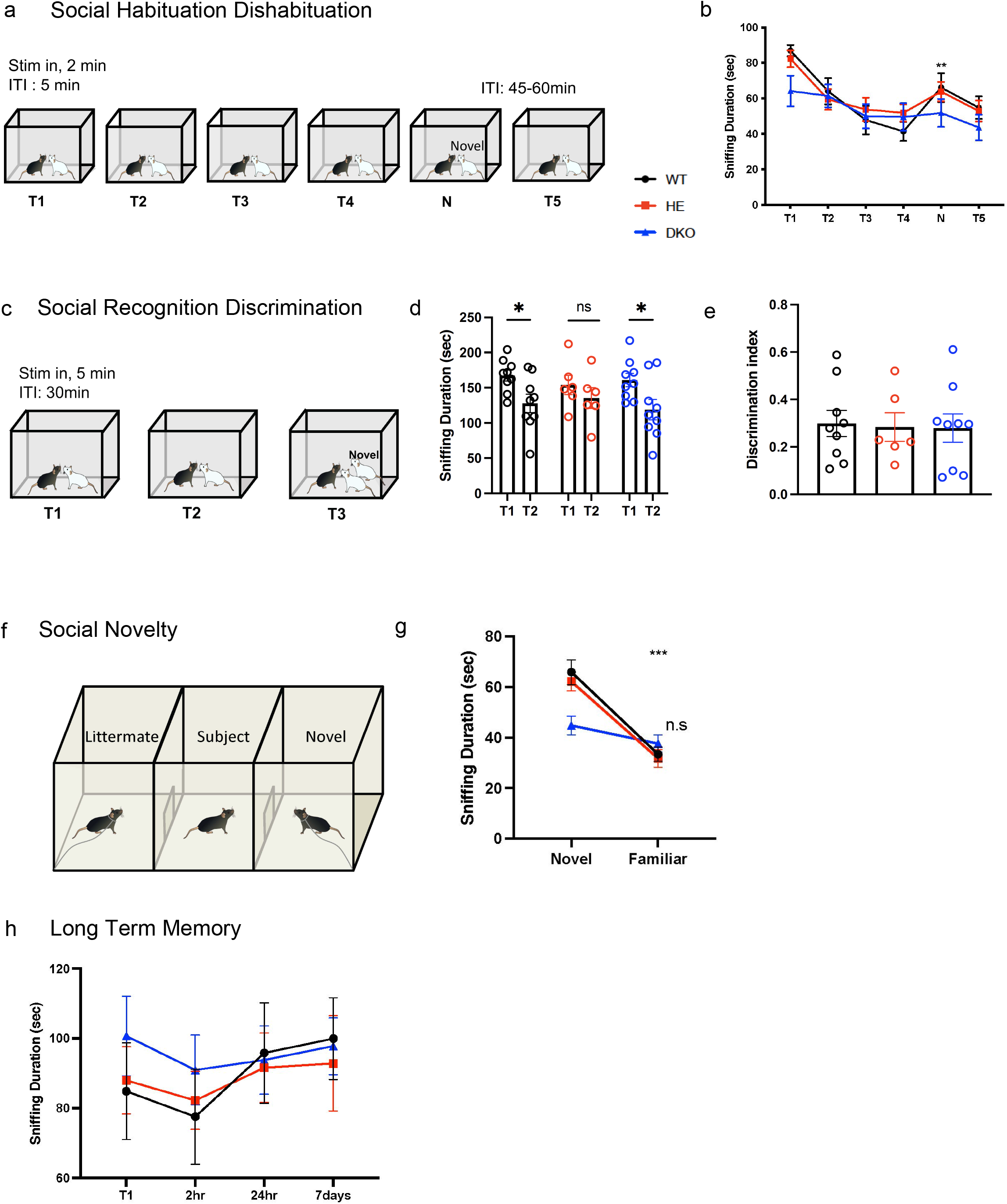
Social memory is impaired following removal of Oxtr or both Oxtr and Avpr1b from CA2 pyramidal neurons. a. Direct social habituation dishabituation test. Trial 1: a subject mouse was presented with a novel Ovx female for 2 minutes. The novel mouse was then removed from the cage for 5 minutes and reintroduced in trials 2-4. In the following trial (N), a novel Ovx female was introduced and finally, after an hour interval the familiar female is reintroduced. b. While the WT group demonstrated habituation to the familiar female and increased sniffing of the novel one, the HE (*Oxtr^-/-^/Avpr1b^+/Cre^*) and DKO (*Oxtr^-/-^/Avpr1b^Cre/Cre^*) mice did not. The HE group habituated to the familiar female but failed to detect the novelty of the female presented in the fifth trial (N) and the DKO mice failed to both habituate and detect the novelty across trials. Two way RM ANOVA of Genotype X Trials, revealed main effect of trials F(4.486,130.1)=18.41 P<0.0001. Tukey’s multiple comparison within genotype showed the significant decrease in sniffing across trials of habituation in WT and HE *P*<0.0005). c. Direct social recognition discrimination test. Trial 1: a subject mouse was presented with a novel Ovx female for 5 minutes. The novel mouse was then removed from the cage for 30 minutes and reintroduced. Following a second interval of 30 minutes, the same female together with a novel one were introduced, for a discrimination trial (T3).d. HE mice had impaired recognition, while the two other genotypes performed the test. Two-way RM ANOVA of Genotype X Trials, revealed a main effect of trial (F(1,21)=13.93;P<0.0012), with Sidak’s multiple comparisons within genotype showing significant decrease for WT and DKO groups (P<0.03). e. There was no significant difference across groups in the discrimination trial (ANOVA between Genotypes F(2,21)=1.287, *P*=0.2971) f. Social novelty test in a three-chamber apparatus. Subject mouse was presented with a littermate on one side and a novel male on the other and allowed 5 minutes of exploration. g. DKO mice failed to detect novelty, indicated by similar sniffing duration of the familiar and novel stimuli. Two-way RM ANOVA of Genotype X Stimulus, revealed main effect of stimulus F(1,41)=63.12;*P*<0.0001 and an interaction F(2,41)7.950;P=0.0012, with Sidak’s multiple comparisons within genotype showing a significant longer sniffing of the novel mouse in the WT and HE mice (P<0.0001). h. Direct social recognition test along growing intervals, in which the subject mouse was repeatedly presented the same ovariectomized female in a clean cage resulted in no significant changes across the genotypes. ITI, inter-trial interval. Results are means ± s.e.m.

The loss of Oxtr (HE), but not both Oxt and Avpr1b receptors (DKO), resulted in impaired ability of male mice to recognize the familiar female in the second trial of the social recognition test (Fig. 2c-d). The DKO mice were able to recognize the familiar female when presented for a longer duration (5 min) and with a longer interval (30 min). Furthermore, there was no significant difference across all tested groups in the discrimination trial (Fig. 2e). These results suggest that all 3 genotypes of mice were able to discriminate between the familiar and novel females when presented simultaneously and with full access. Yet, in a separate social novelty test in which mice had to discriminate between a male littermate and a novel male mouse (Fig. 1f, g), the DKO group (*Oxtr^-/-^/Avpr1b^Cre/cre^*) failed. The mice did not detect the novelty of the mouse presented and spent similar time sniffing it compared to the littermate (Fig. 1h).

Finally, in order to investigate the role of the receptors in long-term social memory, we tested the mice with longer intervals of up to 7 days from the initial presentation. There were no significant differences between groups along all trials (Fig. 1h). All mice, regardless of genotype, failed to recognize the familiar female when presented to them 2 hours, 24 hours or 7 days after the initial presentation. The lack of memory in these retention times is in agreement with our previous observations^2^ and others^2, 25^.

### Conditional knockout of *Oxtr* or both *Oxtr* and *Avpr1b* in CA2 pyramidal neurons impairs non-social novelty detection

While acute removal of Oxtr in CA2/aCA3 did not affect non-social recognition nor discrimination in a previous study^23^, impaired approach to non-social novelty was observed following Oxtr pharmacological or genetic inactivation^26^. In this study, preference for novel non-social stimulus was rescued by habituation to either context or the object. Impaired object novelty detection was also observed in Oxytocin receptor mutant zebrafish, further demonstrating the receptors’ involvement in a more general memory recognition process of familiar versus novel entities^27^.

Here, all groups habituated to the repeatedly presented object, yet the HE and DKO groups had impaired novelty detection of the object presented in trial N (Fig. 3 a-b). Moreover, the HE mice failed to recognize the familiar object when presented again one hour later (T5). Thus, our data suggest the Oxt receptor in the pyramidal neurons of CA2 is involved with the process of non-social novelty detection.

**Figure 3.**
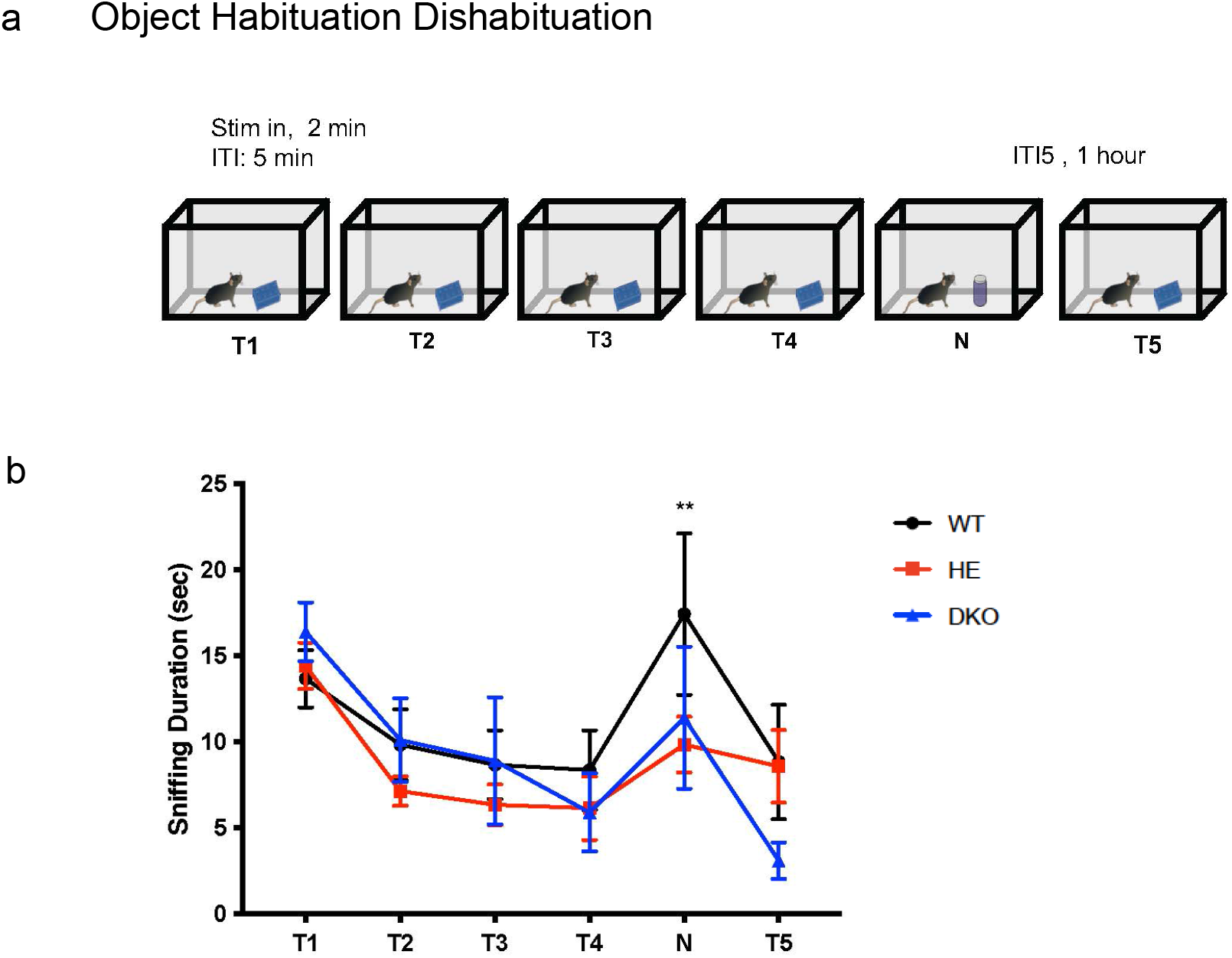
Object Novelty detection, but not recognition, is impaired by removal of Oxtr alone or both Oxtr and Avpr1b from CA2 pyramidal cells. a. Direct object habituation-dishabituation test. Trial 1: an object is presented to the subject mouse to explore for 2 minutes. The object is then removed from the cage for 5 minutes and reintroduced in trials 2-4. In the fifth trial (N), a new object is presented and finally after a prolonged interval of one hour, the familiar object is presented once again. b. The groups differ significantly (Two-way ANOVA: Genotype X Trial F(10,145)=1.995, *P*<0.04; Trial F(5,146)=13.27, *P*<0.0001). Mice lacking Oxtr (HE) or Oxtr and Avpr1b (DKO) in the CA2 pyramidal neurons did not detect the noveltly of the object in trial N. ITI, inter-trial interval. Results are means ± s.e.m.

### Synaptic transmission between CA3 and CA2 is enhanced by conditional knockout of both *Oxtr* and *Avpr1b*

CA2 is centrally located and well connected within the hippocampus, serving as an intra-hippocampal hub controlling hippocampal excitability. Pyramidal cells in CA2 receive direct excitatory input from CA3^28, 29^ as well as project to area CA3, where they recruit inhibition^29^. They also make strong excitatory synapses with CA1^28^. Furthermore, Oxt exerts an early and glutamatergic-specific ‘priming’ effect on synaptic development in hippocampal neurons in culture, through the Oxt receptor^30^.

To examine how the loss of the Oxtr and Avpr1b impacts hippocampal physiology, we recorded evoked fEPSPs in the SR of the dorsal CA2 or in the SO of the dorsal CA1 while stimulating terminals of CA3 or CA2, respectively (Fig.4a). We observed enhanced synaptic transmission between CA3 and CA2 in the DKO mice (*Oxtr^-/-^ Avpr1b*^Cre/Cre^) but no change in HE mice (*Oxtr^-/-^Avpr1b*^+/Cre^) (Fig. 4b).

**Figure 4.**
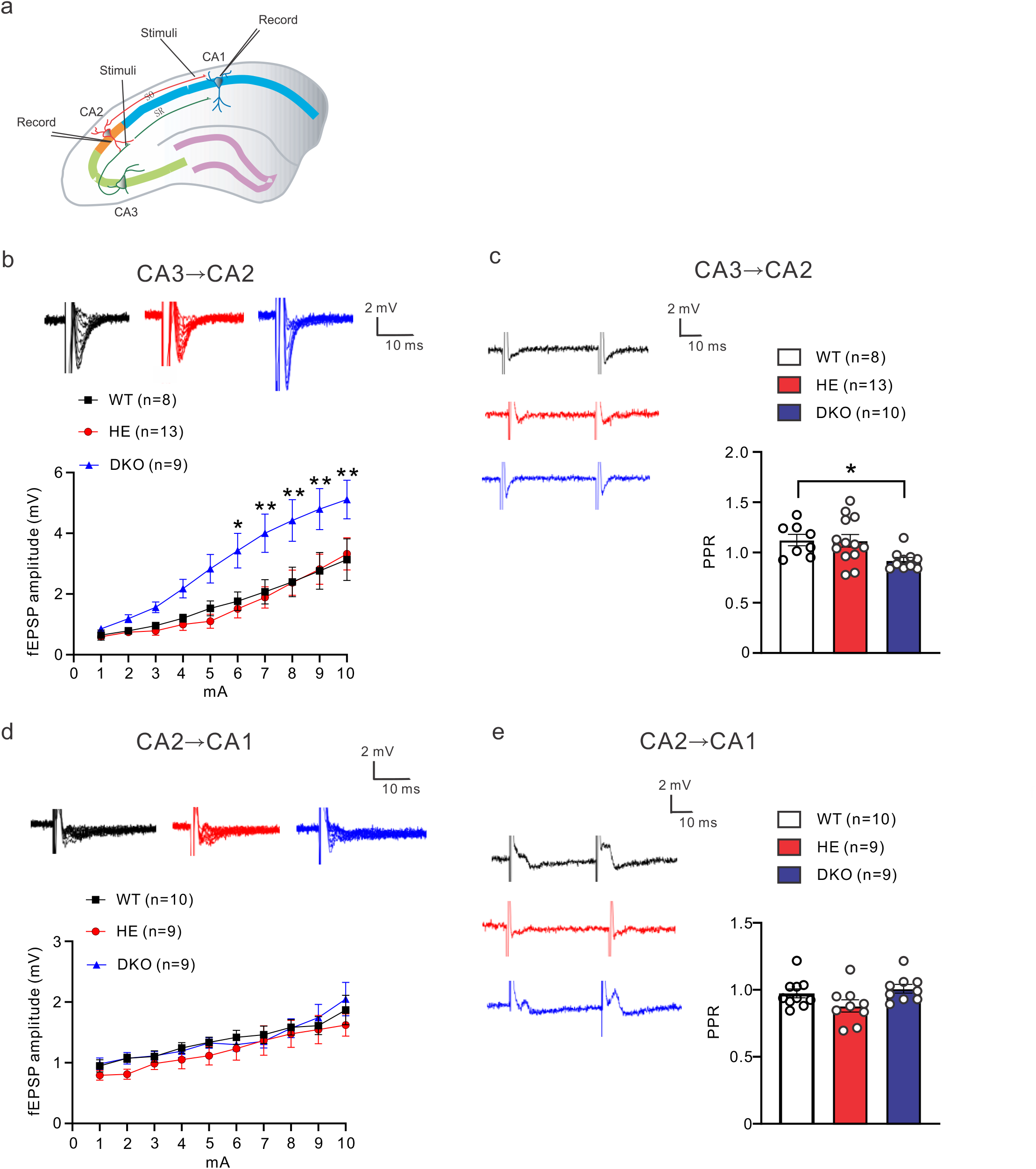
Synaptic transmission between CA3 and CA2 is enhanced in the DKO. a. Strategic illustration for electrophysiological recordings. b. fEPSP amplitude in CA2 pyramidal neurons was increased in the DKO mice, following stimulation of CA3 terminals. Two-way ANOVA of Genotype X Stimulus intensity (Amplitude) revealed a main effect in both Genotype F(2,270)=38.92, P<0.0001 and Stimulus intensity F(9,270)=22.71, P<0.0001, with Tukey’s multiple comparison further demonstrating higher fEPSPs amplitudes in the DKO group in stimulus intensities of 5mA and above. c. Paired -pulse ratios (PPR) were compared using ANOVA, showing a main effect across genotypes (F(2,28)=4.260, P=0.024), with Dunnett’s multiple comparisons further identifying a lower PPR in mice lacking both receptors (DKO;*P*<0.05). d. fEPSP amplitude in CA1 pyramidal neurons was unchanged in the HE or DKO mice, following stimulation of CA2 terminals. e. PPR in CA1 were similar across genotypes. n indicates the number of brain slices from 3 animals.

In order to identify changes in the population of presynaptic terminals, we tested short-term plasticity with a paired-pulse protocol within a short inter-pulse interval of 50ms^31^. Paired-pulse ratios (PPR) were lower in DKO mice (Fig. 4c). This indicates that short-term plasticity is affected by the loss of Oxtr and Avpr1b from the pyramidal neurons in CA2 and suggests that the presynaptic release probability of CA3 neurons is increased. We then tested whether the loss of the Oxtr and/or Avpr1b alters presynaptic output of CA2 onto its major downstream target CA1. The synaptic transmission between CA2 and CA1 was unaffected by the loss of either the Oxtr or both Oxtr and Avpr1b (Fig. 4d-e). These results suggest the co-expression of Oxtr and Avpr1b is critical for maintaining synaptic transmission between CA3 and CA2.

### Spontaneous activity is changed in CA2 stratum oriens by conditional knockout of both Oxtr and Avpr1b

To further test whether the loss of the receptors from the pyramidal neurons of CA2 has any functional impact, we analyzed spontaneous local field potentials (sLFPs) recorded in both SR (Fig. 5 a, c, e) and SO (Fig. 5 b, d, f). We observed increased power in the delta band (0.5-4Hz) in the SO layer following the loss of both Oxtr and Avpr1b (Fig. 5f), suggesting a selective shift in the excitatory/inhibitory balance within the apical dendrites layer and modulation of the network rhythm.

**Figure 5.**
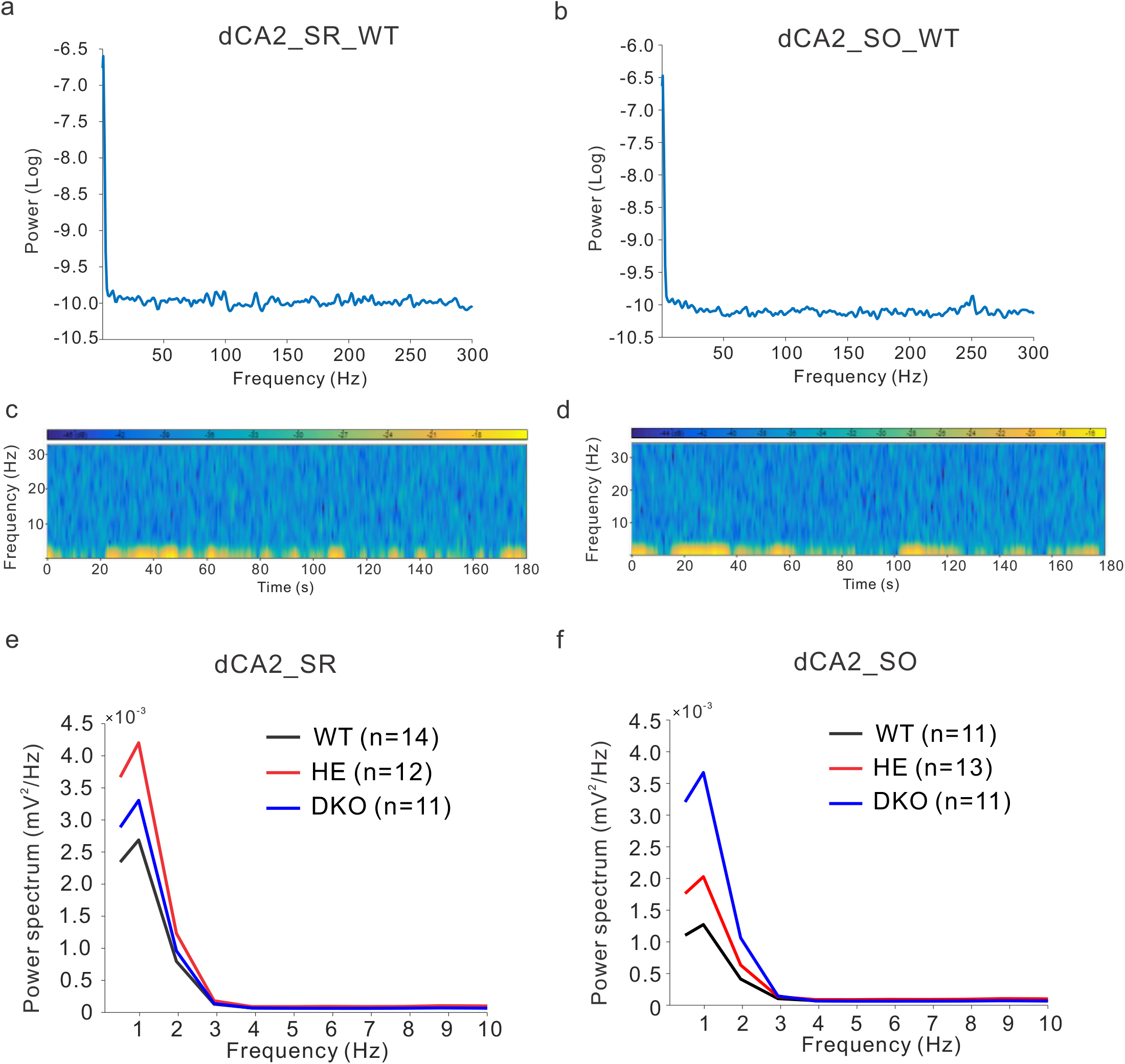
The spontaneous local field potentials in CA2 SO are increased by the removal of both Oxt and Avp1b receptors. (a, b) An example of power spectrum in CA2 region shown from a WT animal. (c, d) An example of time course power spectrum in CA2 SR (c) or SO (d) estimated from a WT animal. (e) Power spectrum analysis in the SR shows no difference between groups. (f). Power spectrum analysis in SO shows increased power in the delta band (0.5-4Hz), for the DKO. Two way ANOVA across frequencies F(10,352)=7.706, *P*<0.0001. Tuley’s mulitple comparisons show DKO significantly higher than WT in 0.5 Hz (P<0.035) and 1Hz (P< 0.003).

### Acute blockade of Oxtr and Avpr1b decreases synaptic transmission between CA3 and CA2

We and others have previously described CA2 induced excitatory potentiation by Oxtr and Avpr1b agonists^17, 32^. Here, we tested the synaptic transmission between CA3 and CA2 in the presence of the Avpr1b antagonist SSR149415 (Fig. 6a), Oxtr antagonist L-371257 (Fig. 6c), or both (Fig.6e). In all conditions, as expected, we observed decreased synaptic transmission. Paired-pulse ratios remained unchanged upon exposure to the antagonist (Fig. 6 b, d, f).

**Figure 6.**
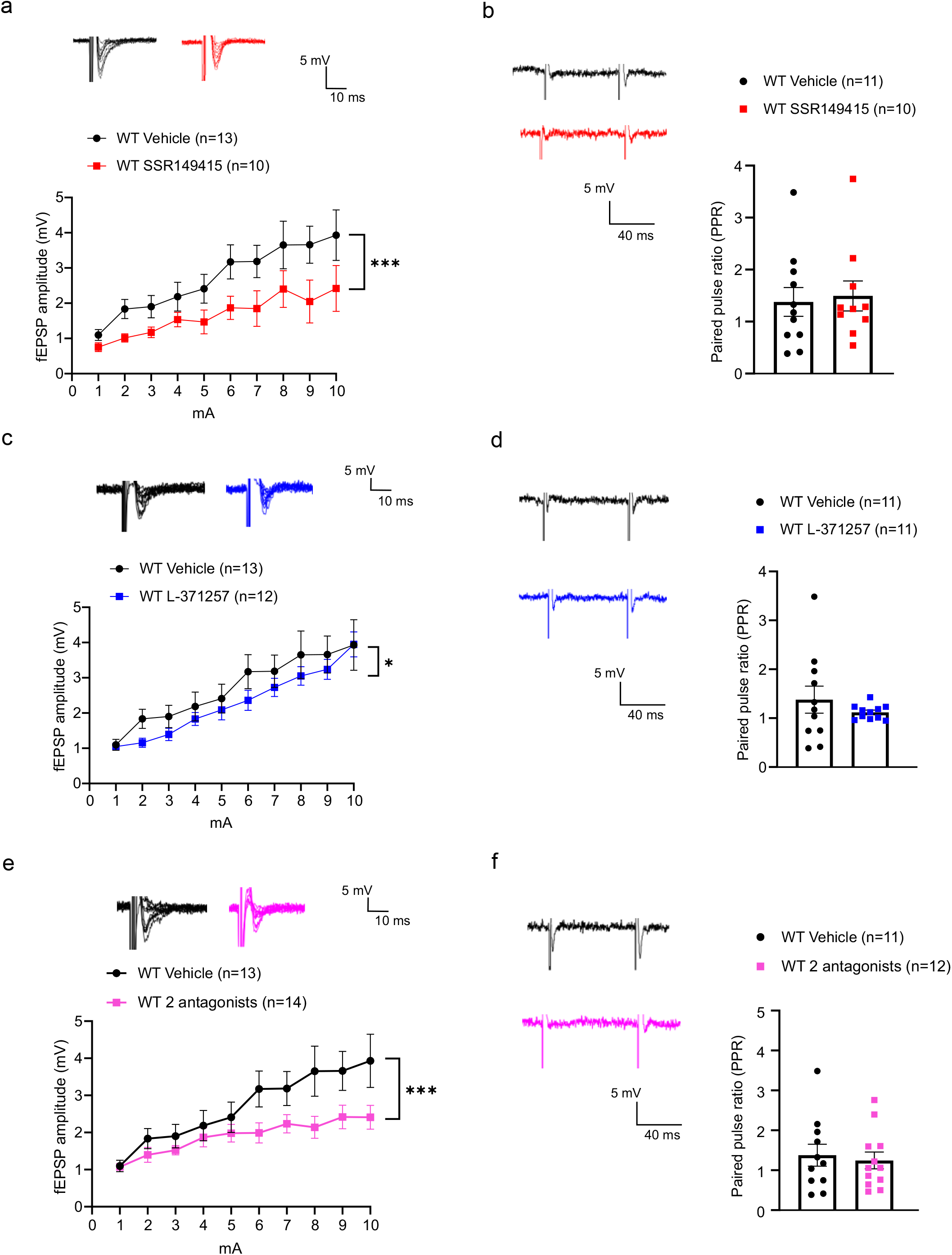
Pharmacological blockade of the receptors reduces activity in dCA2. a. SSR149415 (Avpr1b antagonist) decreased fEPSP amplitude in CA2 pyramidal neurons. Two-way ANOVA demonstrated effect of treatment F(1,210)26.43, P<0.0001. b. No change was observed in the PPR. c. L-371257 (Oxtr antagonist) decreased fEPSP amplitude in CA2 pyramidal neurons. Two-way ANOVA demonstrated effect of treatment F(1,230)=6.974, P<0.02 d. No change was observed in the PPR. e. Combining both SSR149415T and L-371257 antagonists decreased fEPSP amplitude in CA2 pyramidal neurons. Two-way ANOVA demonstrated effect of treatment F(1,250)=23.42, P<0.0001. f. No change was observed in the PPR. n indicates the number of brain slices from 3 animals.

## Discussion

Avp and Oxt are important neuromodulators implicated in many central processes with rapid behavioral effects. While their receptors are distributed across multiple brain regions, the specific Avp receptor, Avpr1b, is colocalized with Oxtr in the pyramidal neurons of the hippocampal CA2 subfield^4, 33, 34^. Although the chemical structures of Avp and Oxt as well as their receptors are similar, they can have distinct or even opposing roles in the regulation of behavior^18^. Yet, cross-talk among the receptors allows them to elicit behavioral changes, even in the absence of either one^18^. While considerable progress has been made, many gaps remain in our understanding of the receptors’ neurophysiological roles in mediating social memory and behaviors.

Previous studies targeting the Oxtr in the CA2 area have described different observations^11, 23^. Discrepancies may arise from methodological differences. Furthermore, given the distribution profile of Oxtr in the hippocampus^35^, acute or chronic viral targeting of the receptor cannot exclude the CA3 subfield. Also, Oxtr is expressed in a development-dependent pattern and its expression levels during a unique developmental window have been suggested to be involved in shaping individual’s behavioral traits^36^. Thus, removal or inactivation of Oxtr in different time windows during development may also be the reason for the different observations across rodent studies. Our tissue-specific approach allowed a developmental point of view on the physiological significance of Oxtr in the pyramidal neurons of CA2. Our data indicates Oxtr actions in the cells are necessary for efficient processing of social and contextual information. Similar to its effect in the medial amygdala^37^, Oxtr actions in CA2 are selective to female stimulus. Previously, we described a failure to recognize an individual female mouse following conditional removal of Oxtr in the forebrain^19^. A later study looking at Ca2+/calmodulin-dependent protein kinase IIα (Camk2α) distribution in mice brain^38^ suggested that the forebrain conditional removal in excitatory neurons included all hippocampal subfields and layers. The current observations of impaired recognition together with their intact ability to discriminate between simultaneously presented females, further delineate the CA2-specific role of Oxtr in individual recognition of females.

Despite the behavioral phenotype, we did not observe changes in spontaneous activity or synaptic transmission between the CA3 and CA2 or CA2 and CA1 following the removal of Oxtr alone. This stands in agreement with a previous publication describing Oxtr-mediated excitatory effects in pyramidal neurons of CA2 with little direct influence on presynaptic terminals of CA2 in CA1^32^. Furthermore, while the CA1 region is highly reactive to novel environmental stimuli^39^ and the presence of a conspecific^40^, no major changes were observed at functional, morphological or biochemical measures in the CA1 of *Oxtr* KO mice^26^, despite an impaired approach to novelty. Further investigation of inhibitory populations and possibly extrahippocampal synaptic transmission may offer more insight into the underlying neuronal changes following the Oxtr removal.

In an editorial comment, Young and Flanagan-Cato^41^ suggest the potential functional significance of cross-talk between Oxt and Avp and their receptors. Restricted removal of both receptors from the pyramidal neurons of CA2 allowed us to investigate the functional significance of the cross-talk in this region. Previously, we showed that the life-long absence of Avpr1b resulted in impaired social recognition in both our knockout^6^ and knockin^5^ lines. We predicted similar, if not greater, impairments with the removal of both Oxtr and Avpr1b. As expected, the loss of both receptors greatly impacted the mice ability to perform in the habituation-dishabituation test. They were unable to recognize the familiar or to detect the novelty of the females presented with a short interval of 5 min. Surprisingly, they did recognize a familiar female after a longer interval of 30 min as well as discriminated between her and a novel one when presented simultaneously. In a separate test, male mice could not discriminate between a male littermate and a novel male. This behavioral phenotype indicates the modulatory effect of the neuropeptides is disrupted and the receptors are necessary for normal processing of social cues. However, the ability to perform the social recognition discrimination test with females suggest a compensatory mechanism, either innate and/or developmental.

Genetic compensation in response to gene knockout is a common phenomenon in order to maintain cellular wellness^42^. Upregulation of related genes and rewiring of the network following the loss of function of the receptors is of considerable interest for future studies. Here, compensation is supported by the observation of enhanced synaptic transmission between CA3 and CA2, indicated by greater amplitude of the fEPSPs, which reflects a summation of excitatory postsynaptic responses evoked by the stimulation of the afferent fibers. The decreased paired-pulse ratio implies a presynaptic change in CA3, with a higher probability of basal transmitter release^43^. Moreover, the increased power of delta band we observed in SO of the CA2 depicts lower excitability and synchronization. Spontaneous rhythmic field potentials in the delta band (0.5-4Hz) have been described in isolated hippocampal slices *in vitro* and are thought to arise from the CA3^44^. This rhythmic activity is dependent on intrinsic cellular properties and the connectivity and strength of both excitatory and inhibitory synapses^45, 46^.

Regulation of excitatory and inhibitory inputs plays a significant role in the dynamic of neuronal networks and is necessary for proper function, whereas abnormal balance is thought to underlie various neurodevelopmental disorders^47^. Restricted chronic silencing of the pyramidal cells in CA2 previously revealed a key role of the region in establishing the dynamic of excitation/inhibition balance required for normal action of the hippocampal formation, specifically through a mechanism of feed forward inhibition in CA3^48^. Our data suggest that Oxtr and Avpr1b co-expression in the pyramidal neurons of CA2 is necessary for the feed forward inhibition of CA3 and the establishment of excitation/inhibition balance.

The high degree of preservation of Oxt/Avp systems across mammalian evolution and the heritability of social behavior in humans^49, 50^, has led to many studies in both animal models and humans. Dysfunctions in these systems have been linked to psychiatric disorders such as autism, anxiety and schizophrenia^50^. The last^50^ also suggests that imaging genetics evidence in human support a model in which Oxtr and Avpr1a variants mediate their associated increased disease risk by modulating the limbic circuit. Additionally, increased delta power has been described in the temporal lobe of schizophrenic patients^51^. Thus, as suggested by Song and Albers^18^, understanding the cross-talk between the neuropeptides and their receptors has the potential for substantial translational relevance that could lead to important clinical breakthroughs.

## Acknowledgments

This research was supported by the intramural research program of the NIMH (ZIA-MH-002498-24).

